# Host-parasite interaction explains variation in prevalence of avian haemosporidians at the community level

**DOI:** 10.1101/432260

**Authors:** L. Garcia-Longoria, A. Marzal, F. de Lope, L. Z. Garamszegi

## Abstract

Parasites are a selective force that shape host community structure and dynamics, but host communities can also influence parasitism. Understanding the dual nature from host-parasite interactions can be facilitated by quantifying the variation in parasite prevalence (i.e. the proportion of infected host individuals in a population) among host species and then comparing that variation to other ecological factors that are known to also shape host communities. Avian haemosporidian parasites (e.g. *Plasmodium* and *Haemoproteus*) are abundant and widespread representing an excellent model for the study of host-parasite interactions. Several geographic and environmental factors have been suggested to determine prevalence of avian haemosporidians in bird communities. However, much remains to be known regarding whether host and parasite traits, represented by phylogenetic distances among species and degree of specialization in host-parasite relationships, can influence parasite prevalence. The aims of this study were to analyze factors affecting prevalence in a bird community and to test whether the degree of parasite specialization on their hosts is determined by host traits. Our statistical analyses suggest that prevalence is mainly determined by the interaction between host species and parasite lineages where tolerance and/or susceptibility to parasites plays an essential role. Additionally, we found that although some of the parasite lineages infected a low number of bird species, the species they infected were distantly related and therefore the parasites themselves should not be considered typical host specialists. Prevalence was higher for generalist than for specialist parasites in some, but not all, host species. These results suggest that prevalence mainly results from the interaction between host immune defences and parasite exploitation strategies wherein the result of an association between particular parasite lineages and particular host species is idiosyncratic.

## INTRODUCTION

Parasites have been suggested as a selective force since they might shape host community dynamics [1], alter interspecific competition and influence energy flow [2]. An essential trait in studies of host-parasite interactions is prevalence (i.e. the proportion of individuals infected by a parasite or pathogen in a population at one point in time) [3]. The importance of prevalence is reflected in the amount of studies showing how major ecological factors shaped its intra- and interspecific variation [4,5]. In bird communities, geographic areas, environmental issues or host species/population have been suggested as factors determining prevalence [6]. However, more studies are still needed in order to understand mainly factors affecting parasite prevalence at the community level.

Avian *Plasmodium* and *Haemoproteus* spp. represent a well-studied host-parasite system. These parasites are vector-transmitted organisms that can cause host mortality or morbidity during the acute phase of infection impacting the life histories of their hosts [7]. Their life-cycle is complex, involving sexual stages in their dipteran vectors and asexual stages in their vertebrate hosts [7]. Prevalence of these parasites may be affected by different factors concerning the vector, hosts and the parasites themselves [7]. Some authors have focused their attention on the possibility that host traits might determine prevalence of these parasites [8–10]. For instance, blood parasite susceptibility of both bird individual [11] and bird species [12,13] have been recently suggested as factors affecting prevalence. In this sense, some bird species may have developed both tolerance and resistance mechanisms [14] underlying the importance of bird species in prevalence studies.

In addition to characteristics of hosts, characteristics of parasites themselves can affect their prevalence in wild communities. Haemosporidians parasites present high plasticity and versatility reflected in the number of parasite lineages found among bird species. Thus, within haemosporidian parasites there are more than 3000 parasite lineages [15] infecting more that 1500 bird species. Each of these lineages may have diverse virulence [16] which could result in an array of different negative effects inside the hosts. The plasticity that malaria lineages may have [17,18] allow these parasites to fully exploit hosts reacting even to changes in the physiological state of the host or the environment [19]. Thus, prevalence displayed by parasite lineage in a determined host species or individual is simply the interaction between host immune defences (susceptibility or tolerance) and parasite strategy (virulence). Additionally, malaria parasites must find the correct host in order to complete its life cycle and achieve a high transmission rate [3,20]. To find the precise host is crucial for specialist parasites that need a particular bird species unlike generalist parasites that infect a wider range of bird species [21]. However, and despite the large number of studies focused on haemosporidian parasites, there are a scarce number of them dealing with factors affecting prevalence in wild communities from the perspectives of both the host and parasites.

*Plasmodium* and *Haemoproteus* spp. might be detected in several host species that can be distantly or closely related to each other [22–24]. Mechanisms allowing parasite to switch host is a central issue nowadays in avian malaria studies [25,26]. Host switching mechanism is a strategy that malaria parasites may use, in order to infect larger number of bird species [27,28], where closely parasite lineages may be detected in bird species with similar immune defences [29–31]. Thus, parasite lineages infecting close related bird species may exhibit the same exploitation strategy for avoiding host defences [32]. Conversely, more generalist parasites would be able to infect a broad range of bird species that might be distantly related [33]. Analysing phylogenetic distances between bird species sharing the same parasite lineages become essential in community studies in order to determine parasite strategies and how specialist and generalist parasites exploit different host species. However, little is know about how haemosporidian parasite lineages infect bird host species depending on the host phylogenetic distance.

The fact that haemosporidian parasites need a vector for completing their cycle life extends the number of elements affecting prevalence of these parasites. Although some studies have shown limited or no relationship between climate and prevalence [34], some have showed that vector populations might increase their number due to climate change and, therefore, vector-borne infectious diseases may increase every year [35]. Vector availability might also change among seasons, thus, during spring the amount of vectors in the wild increase [36] raising the probabilities to become infected [37]. During this season the probability of becoming infected increases significantly not only because of vector availability but also because of the secretion of sexual hormones that may alter immune system and, therefore, facilitate the entrance of blood parasite in the vertebrate host [38]. These studies emphasize the importance of including both year and season of sampling for studies focused in prevalence of haemosporidians prevalence when analysing communities.

Taken as a whole, studies focused on prevalence at the community level have analysed certain factors affecting the presence of prevalence separately. However, to date there is no study dealing with the main factors (i.e. host species, parasite lineage and vector conditions) that might affect the presence of haemosporidians parasites or with the parasite strategy depending on the host phylogenetic distance. Hence, the aims of this study are (i) to analyse factors affecting prevalence of haemosporidian parasites in a wild bird community such as phylogenetic distances, parasite phylogeny, host ID, parasite lineage or seasonal effects and (ii) compare the phylogenetic distances among bird species infected with the same parasite lineage in order to determine whether the parasite infects phylogenetic close host species. For this purpose, factors such as year, day of sampling, bird species, parasite lineage, season of sampling, host-parasite interaction and phylogenetic relationships among avian hosts were included in the analyses and related with prevalence of haemosporidian parasites.

## MATERIAL AND METHODS

### Data collection and database

The study was carried out from February to October during a 9-year period (2002–2010) in the surroundings of Badajoz (SW Spain) (38°52’N, 705’W). Birds were collected always during the sunrise and during the early hours of the morning. Using a mist-net system, we captured a total of 815 individuals from 21 bird species belonging to nine different families. We obtained one microcapillary of blood (70μl) from the brachial vein of each individual and stored it in 500 μl of 96% ethanol until analysis.

Methods were approved by Institutional Commision of Bioethics of Univ. of Extremadura (CBUE 49/2011)

### Prevalence and genetic detection of parasite lineages

DNA from the avian blood samples were extracted in the laboratory using a standard chloroform ⁄ isoamylalcohol method [39]. Diluted genomic DNA (25 ng/μL) was used as a template in a polymerase chain reaction (PCR) assay for detection of the parasites using the nested-PCR protocols described [40]. The amplification was evaluated by running 2.5 μL of the final PCR on a 2% agarose gel. All PCR experiments contained one negative control for every eight samples. In the very few cases of negative controls showing signs of amplification (never more than faint bands in agarose gels), the whole PCR-batch was run again to make sure that all positives were true. All samples with positive amplification were sequenced directly using procedures described [41]. The obtained sequences of 478 bp of the cyt b were edited, aligned and compared in a sequence identity matrix using the program BioEdit [42]. Parasites with sequences differing by one nucleotide substitution were considered to represent evolutionary independent lineages [43]. Five new sequences were deposited in GenBank under the accession numbers JQ749720 – JQ749724.

### Phylogenetic analyses and statistical procedures

For our statistical models (see below) we created two different phylogenetic trees (supporting information Figs S3 and S4) to control for the common descent of different parties: one for the host of bird species and one for the parasite lineages. The first one relied on the 1000 trees generated by the birdtree.org website [44], from which a consensus tree was created for the sampled 21 species by using Geneious v5.4 [45]. The second tree relied the sequences of the parasite lineages [15] identified in this study. Then, we created a phylogenetic tree using MrBayes 3.1 [46,47]. We used jModelTest 0.1 [48] in order to determine which parasite tree offered the best fit to our data. The burn-in was identified through Tracer 1.2.2 [49]. We sampled 10 million generations at intervals of 1000. Finally we analysed the files generated by Bayesian MCMC runs by MrBayes in Tracer with the objective of confirm whether the parasite tree generated was the most adequate to our analyses.

These phylogenetic trees were entered in the subsequent statistical models sequentially to evaluate the same list of predictors (first, we run the model using the bird phylogenetic tree and then we run the same model but we used the parasite phylogenetic tree). The general strategy for our modelling was to incorporate different factors that might be related to: (i) vector effects (breeding season or non-breeding season, year and date of sampling), (ii) host effects (host species, phylogenetic relatedness), (iii) parasite effects (lineage identity and parasite phylogenetic relatedness), and (iv) host-parasite interaction (the combination of bird species and parasite lineage). Given that our parasite screening method allowed us to detect prevalence (yes or no) at the individual level for each parasite lineage screened, we could also incorporate effects due to host individual identity. We used the Bayesian framework for generalized mixed models (GLMM) incorporating Markov chain Monte Carlo (MCMC) estimation available in the package ‘MCMCglmm’ [50]. Because dependent variable (a given parasite lineage detected or not in a given individual of a given species) has a binomial distribution, we adopted the “categorical” family of distribution. The fixed variables we considered were the date of sampling and the breeding status of individuals (breeding or not), as these factors may affect infection status due to mosquito abundance/activity and susceptibility of hosts during the demanding chick-feeding period [7]. The year of sampling, parasite lineage, bird individual identity, host species and phylogenetic relationships (parasite or host) were used as random factors. Additionally, we also included the interaction between host species and parasite lineage as a random factor [11,51]. Plots were made with the R package ggplot2 (v. 2.1.0) [52]. All the statistical analyses were carried out with the program R v.1.1.383 [53].

We used uninformative priors with a low degree of belief in all parameters. The model was run for 130000 iterations preceded by a burn-in of 30000 iterations, and sampling every 100 iterations to avoid autocorrelation. We evaluated model convergence visually by plotting the chains and checking that they had mixed properly and by plotting the autocorrelation (supporting information Figs S1 and S2). Once both model were finished we rescaled our parameters by estimating the marginal parameter modes using Kernel Density Estimation [50].

We examined phylogenetic relatedness of bird species infected by the same lineages, and determined if species with non-zero prevalence are more closely related to each other than could be expected by chance. The significance of this metric was tested by comparison to a null distribution derived from 999 random permutations among the tips of the phylogenetic tree followed by calculation of the main pairwise distance (MPD). Calculations were carried out with the ses.mpd function of the R package Picante [54]. When running a main pairwise distance calculation different value comes out. Thus, MPDobs = observed mean pair-wise phylogenetic distance between all species pairs infected with the parasite. The mean and standard deviation of MPD in the null distribution were obtained by randomization of species in the phylogenetic distance matrix (taxa.labels method in Picante). Z = (MPDobs – mean MPD of the null distribution)/SD of the null distribution. Negative values of Z indicate greater phylogenetic homogeneity (clustering). P is the probability of drawing an MPD from the null distribution at least as extreme as MPDobs, based on 999 randomizations.

## RESULTS

### Total prevalence and lineages detected

We determined the presence of 26 parasite lineages belonging to the genera *Haemoproteus* (N = 13) and *Plasmodium* (N = 13) in the analysed host species (Table 1). We found that 63.80% of total individuals and 81 % of species were infected by haemosporidian parasites. We detected five new parasite lineages that had not been previously described (Table 2).

**Table 1.**
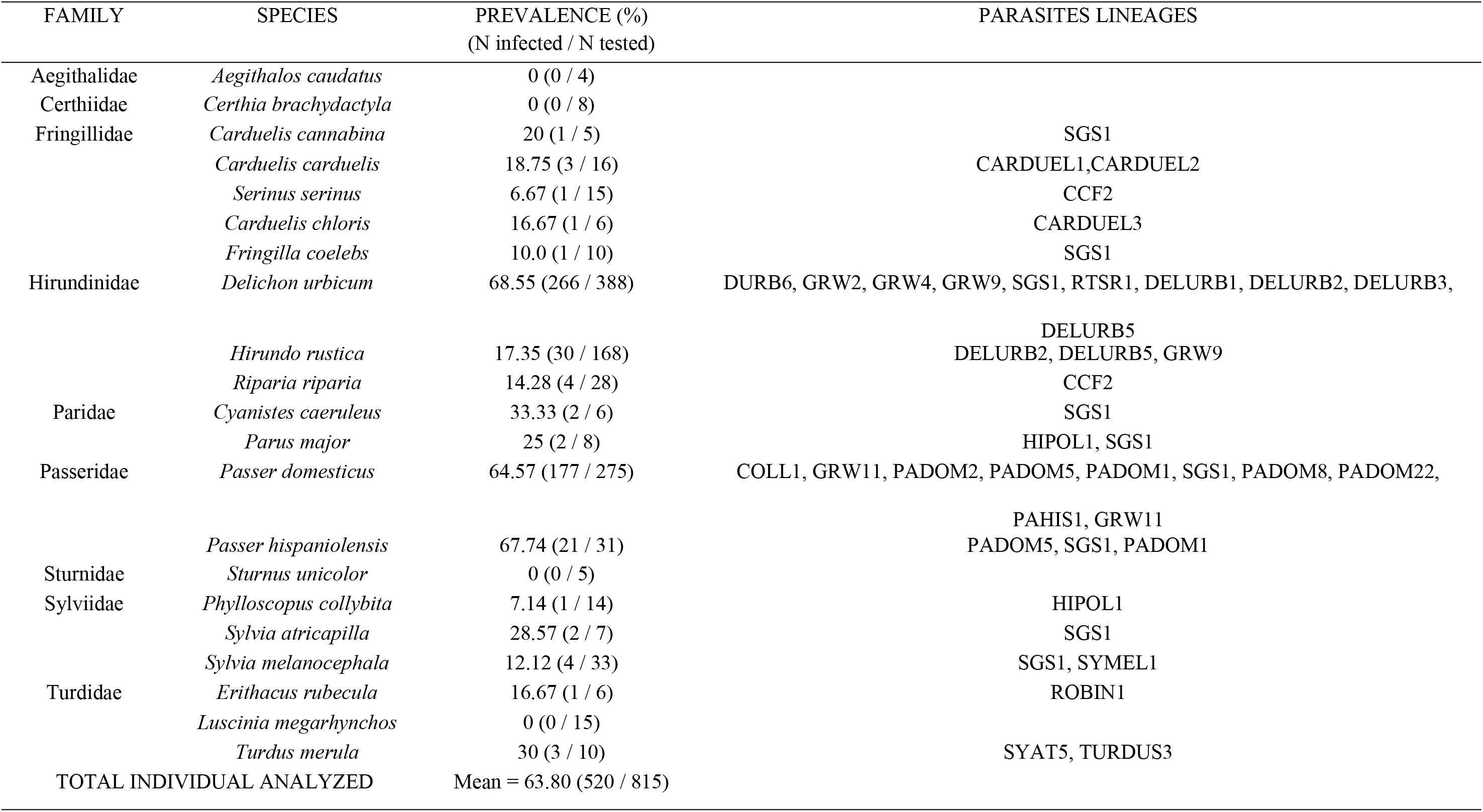
Prevalence and genetic identity of haemosporidian parasite lineages in Passeriform host species analyzed in this study. and parasite lineage found in each host species.

| FAMILY | SPECIES | PREVALENCE (%)<br>(N infected / N tested) | PARASITES LINEAGES |
| --- | --- | --- | --- |
| Aegithalidae | *Aegithalos caudatus* | 0 (0 / 4) |  |
| Certhiidae | *Certhia brachydactyla* | 0 (0 / 8) |  |
| Fringillidae | *Carduelis cannabina* | 20 (1 / 5) | SGS1 |
|  | *Carduelis carduelis* | 18.75 (3 / 16) | CARDUEL1,CARDUEL2 |
|  | *Serinus serinus* | 6.67 (1 / 15) | CCF2 |
|  | *Carduelis chloris* | 16.67 (1 / 6) | CARDUEL3 |
|  | *Fringilla coelebs* | 10.0 (1 / 10) | SGS1 |
| Hirundinidae | *Delichon urbicum* | 68.55 (266 / 388) | DURB6, GRW2, GRW4, GRW9, SGS1, RTSR1, DELURB1, DELURB2, DELURB3,<br>DELURB5 |
|  | *Hirundo rustica* | 17.35 (30 / 168) | DELURB2, DELURB5, GRW9 |
|  | *Riparia riparia* | 14.28 (4 / 28) | CCF2 |
| Paridae | *Cyanistes caeruleus* | 33.33 (2 / 6) | SGS1 |
|  | *Parus major* | 25 (2 / 8) | HIPOL1, SGS1 |
| Passeridae | *Passer domesticus* | 64.57 (177 / 275) | COLL1, GRW11, PADOM2, PADOM5, PADOM1, SGS1, PADOM8, PADOM22,<br>PAHIS1, GRW11 |
|  | *Passer hispaniolensis* | 67.74 (21 / 31) | PADOM5, SGS1, PADOM1 |
| Sturnidae | *Sturnus unicolor* | 0 (0 / 5) |  |
| Sylviidae | *Phylloscopus collybita* | 7.14 (1 / 14) | HIPOL1 |
|  | *Sylvia atricapilla* | 28.57 (2 / 7) | SGS1 |
|  | *Sylvia melanocephala* | 12.12 (4 / 33) | SGS1, SYMEL1 |
| Turdidae | *Erithacus rubecula* | 16.67 (1 / 6) | ROBIN1 |
|  | *Luscinia megarhynchos* | 0 (0 / 15) |  |
|  | *Turdus merula* | 30 (3 / 10) | SYAT5, TURDUS3 |
| TOTAL INDIVIDUAL ANALYZED |  | Mean = 63.80 (520 / 815) |  |

**Table 2.**
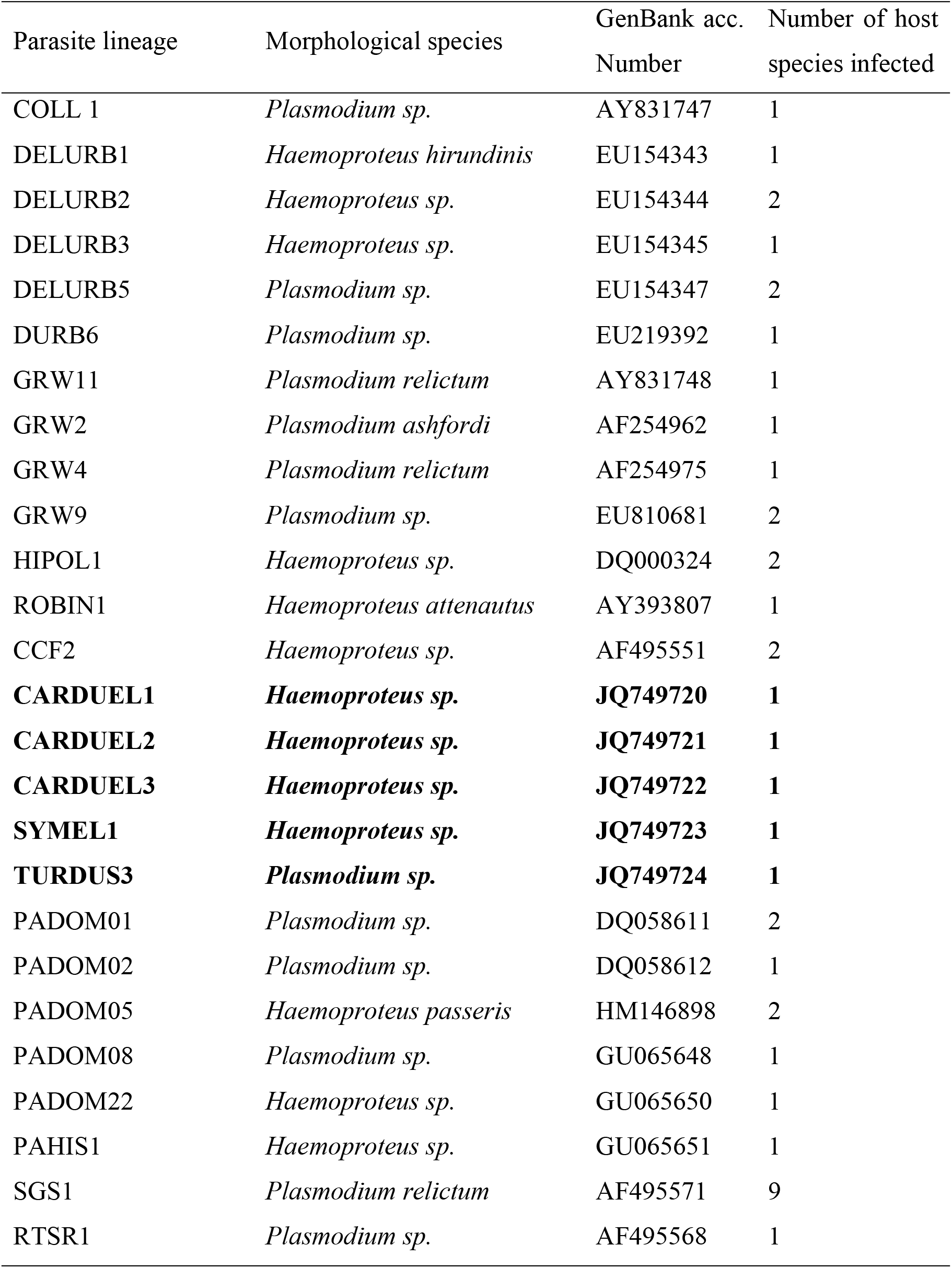
List of parasite lineages found in the current study. New detected parasite lineages are marked in bold.

### Factors affecting prevalence

Our results suggest that bird defences and parasite virulence (i.e. host-parasite interaction) affect infection status stronger than the rest of the variables analysed (Table 3) where this interaction might be the most important factor determining prevalence. Additionally, two more factors appeared to affect variance in prevalence, although on a lesser scale. Thus, our models suggested that breeding season and parasite lineage might affect haemosporidian prevalence.

**Table 3.** Variance of prevalence in relation to fix and random factors. Results of both models are showed. Bolded values represent factors that were statistically significant.

|  | Bird |  |  |  | Parasite |  |  |  |
| --- | --- | --- | --- | --- | --- | --- | --- | --- |
|  | Mean | Lower/upper 95% CI | pMCMC | Rescaled mean | Mean | Lower/upper 95% CI | pMCMC | Rescaled mean |
| *Fixed factors* |  |  |  |  |  |  |  |  |
| Day | -4.055e-03 | -8.653-03 / 7.416e-04 | 0.108 | -0.004 | -3.503e-03 | -8.545e-03 / 7.867e-04 | 0.152 | -0.035 |
| **Breeding season** | **1.825e+03** | **-3.292e-01 / 4.2623+00** | **0.092** | **2.071** | **1.7373** | **-1.212e-01 / 3.842** | **0.074** | **1.775** |
| *Random factors* |  |  |  |  |  |  |  |  |
| Year | 1.616 | 0.243 / 4.364 |  | 0.415 | 1.367 | 0.181 / 3.374 |  | 0.578 |
| **Lineage** | **2.208** | **0.0002 / 4.897** |  | **1.335** | **1.436** | **0.0002 / 3.600** |  | **1.009** |
| Individual | 0.006 | 0.0001 / 0.018 |  | 0.005 | 0.007 | 0.0002 / 0.021 |  | 0.007 |
| Species | 0.126 | 0.0003 / 0.567 |  | 0.007 | 0.166 | 0.0001 / 0.845 |  | 0.006 |
| **Linage\* Species** | **13.26** | **8.318 / 21.97** |  | **7.007** | **11.030** | **6.785 / 15.590** |  | **7.076** |
| Phylo | 0.008 | 0.0002 / 0.028 |  | 0.007 | 0.1629 | 0.0002 / 0.747 |  | 0003 |

### Infected bird species and phylogenetic distances

The random effects for parasite lineage shows that there are considerable differences among parasite lineages in their host exploitation strategies. We found that the most prevalent parasite lineage was Plasmodium relictum SGS1 as it was detected in 9 out 21 host species. However, the prevalence in particular host species was quite variable (see Fig. 1). In contrast, the other lineages seem to be more specialists, at the present study, as they were detected in fewer species.

**Figure 1:**
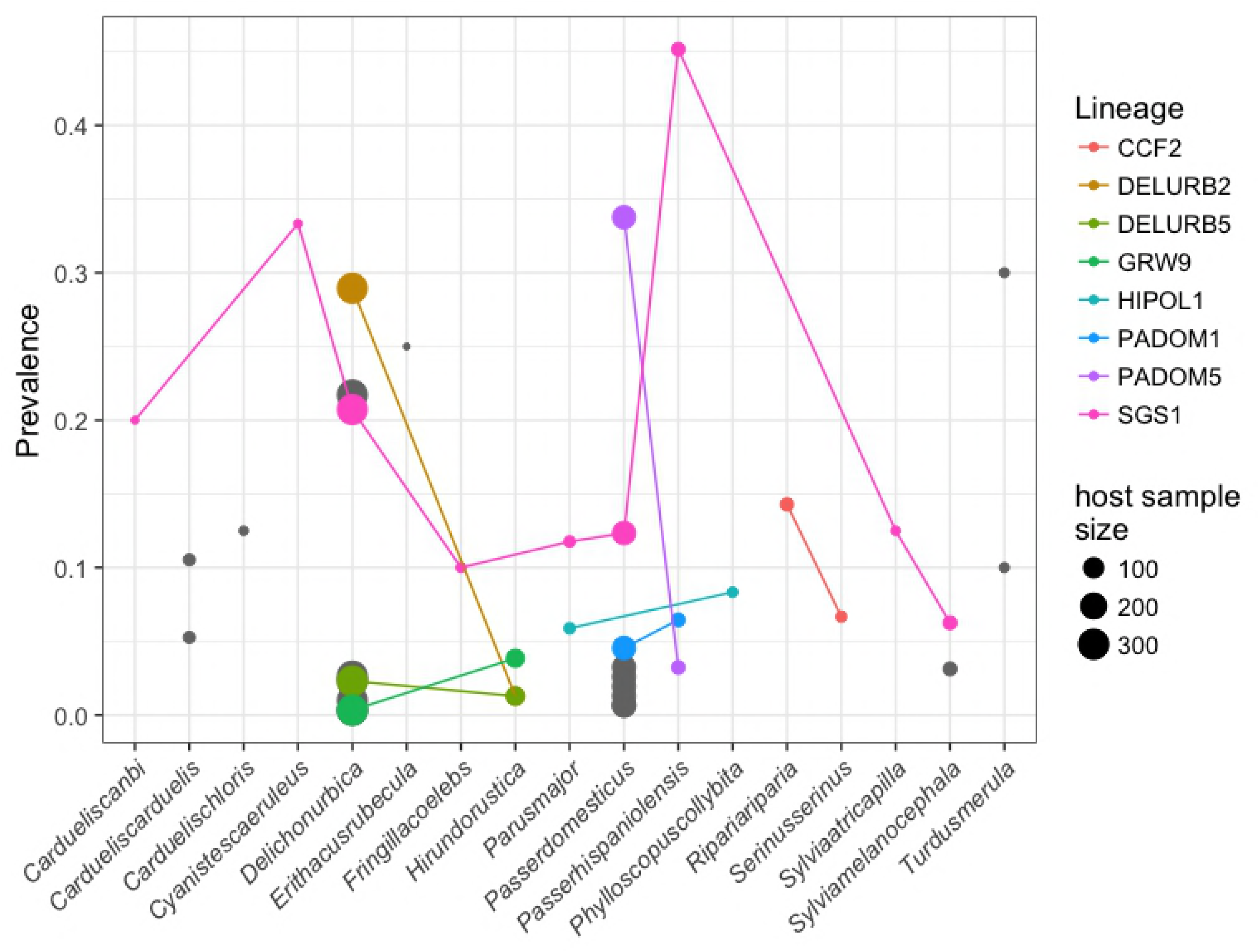
Prevalence of each parasite lineage in every host species found in the community. Bird species with no infection are not present in the figure. Lineages infecting more than one bird species have been colored marked while parasite lineages infected only one host species remain in grey. Noted that depending on the sample size the dot showing the prevalence change its magnitude.

We detected 8 different lineages infecting, at least, two host species (Fig. 1). Thus, we analysed the phylogenetic distance between the hosts species infected by one relative to the average phylogenetic distance between two species on the entire tree. Our results showed that *Haemoproteus* spp. CCF2 lineage, *Plasmodium* spp. HIPOL1, GRW9 and *P. relictum* SGS1 infected non-closely related bird species. Moreover, we detected that *Haemoproteus passeris* PADOM05 lineage; *Haemoproteus* spp. DELURB2 lineage and Plasmodium spp. DELURB5 and PADOM01 lineages tended to infect closely related bird species (Table 4).

**Table 4:** Phylogenetic dispersion of parasite lineages infecting more than one host species. *N* = number of bird species infected. Bolded values represent parasite lineages that were detected in close phylogenetic bird species.

|  | *N* | MPD<sub>obs</sub> | Mean ± SD | *Z* | *P* |
| --- | --- | --- | --- | --- | --- |
| SGS1 *P. relictum* | 9 | 79.976 | 81.829 ± 2.609 | -0.710 | 0.222 |
| **PADOM05 *H. passeris*** | **2** | **5.546** | **82.695±17.199** | **-4.485** | **0.002** |
| CCF2 *Haemoproteus* sp. | 2 | 91.536 | 80.881±18.287 | 0.582 | 0.853 |
| **DELURB2 *Haemoproteus* sp.** | **2** | **33.574** | **82.087±17.088** | **-2.838** | **0.038** |
| **DELURB5 *Plasmodium* sp.** | **2** | **33.574** | **82.111±17.851** | **-2.713** | **0.047** |
| GRW9 *Plasmodium* sp. | 2 | 33.574 | 81.479±18.615 | -2.573 | 0.054 |
| HIPOL1 *Haemoproteus* sp. | 2 | 86.983 | 81.817±17.714 | 0.291 | 0.304 |
| **PADOM1 *Plasmodium* sp.** | **2** | **5.546** | **81.385±18.107** | **-4.188** | **0.003** |

## DISCUSSION

In this study of haemosporidian parasites infecting a community of wild birds breeding/migrating in the same area, we found that detected prevalence is mostly determined by the interaction between host and parasite identity. We also found that variation in prevalence might be affected by breeding season and parasite lineage. Additionally, we showed that *P. relictum* SGS1 is present in most of the bird species but with variable prevalence, while other parasites are more specialist parasites infecting fewer and close related host species as *H. passeris* PADOM05. Next, we discuss these main results in detail.

Virulence of *Plasmodium* and *Haemoproteus* spp. might be highly variable between species and lineages. Some malaria parasites might affect host fitness [55,56] and others might even influence survival [57–59]. Conversely, host bird species and bird individual might also differ in the prevalence displayed due to the tolerance or susceptibility of each organism [11]. Thus, the interaction between parasite lineage and host bird species ultimately determines infection severity and therefore prevalence. Our results support this idea where variation in prevalence was mainly affected by the interaction between parasite lineage and bird host species. In this sense, it has been suggested that avian malaria parasites (*Plasmodium ashfordi*) express a particular host-specific expression pattern depending on the infected bird [11]. This study highlighted the role that the interaction between host immune defence (susceptibility or resistance to parasites) and parasite strategy (virulence) may play in the prevalence displayed. The mechanisms behind this host-parasite interaction are still limited where only some target genes related, for instance, with red blood cell invasion have been deeply analysed [60,61].

With regard to the previous results, prevalence was also affected by parasite lineage. These results emphasize the importance not only of the host-parasite interaction per se but also the parasite exploitation strategy that might vary depending on the bird identity [11]. In this sense, it has been shown that avian malaria parasites might develop a wide range of different strategies in order to avoid host defences. For example, *Plasmodium* spp. can display some evasive mechanisms evading host defences by clonal antigenic switches [62] or even avoid consequences of recognition by the immune system [63]. Additionally, it has been shown that, during mix infections, these parasite might compete with one another for access to the available hosts, thus, some lineages show more prevalence in certain bird species simply because an specific lineage may eliminate other parasite [23]. Hence, parasite lineage might play an essential role determining the prevalence displayed in a host rather the host bird species itself.

Our results also showed that breeding season affected prevalence displayed in the community. Previous studies have shown that, during breeding season prevalence of these parasite significantly increase [64,65] as a consequence of vector availability. The increase of vectors in the community may expand blood parasites among individuals and, therefore, increase prevalence [36]. Our results agree with this idea pointing out a clear connexion between breeding season, vector availability and prevalence of blood parasites.

It has been extensively discussed how specialist and generalist parasites may have evolved and how host species are selected to be infected [22,66,67]. Although specialist parasites are supposed to display higher prevalence and parasitemia on their avian hosts [21], generalist parasite can occasionally cause significant mortalities in native birds. For instance, in Hawaii birds Plasmodium relictum caused the extinction of native species of honeycreepers [68,69] despite being one of the most generalist parasite species. In this sense, it has been suggested that generalist parasites (*P. relictum* GRW4 and SGS1) have the ability to infect a broad range of host species but also display a high prevalence in some of them [70]. Our results agree with this hypothesis, where *P. relictum* SGS1 was detected in most of the bird species infected, and it displayed different mean prevalence depending on the bird species. Moreover, this haemosporidian lineage was detected in unrelated host species highlighting the parasite’s ability to adapt to distantly related host species emphasizing the parasite’s generalist strategy. In the case of P. relictum SGS1 the number of potential host species is 113 [15], where every host individual and species provide a different environment that need to be overcome by the plasticity of parasites [71]. Although the mechanism underlying this plasticity is still unknown, it has been recently shown that the reticulocyte binding proteins, a polymorphic gene family involved in host cell invasion and attributed to host specificity [61], is significantly expanded in P. relictum [72]. This discovery could explain the ability of this parasite species to infect a wide range of avian species and its overwhelming capacity to infect a wide range of host species [72]. Nevertheless, further studies are needed in order to assess mechanisms allowing generalist parasites to be present in a large number of bird species.

Finally, our analysed stated that the number of infected bird species by the same parasite lineage do not determine whether this parasite may be more or less specialist. In this sense, *Haemoproteus* spp. CCF2, HIPOl1 lineages and *Plasmodium* spp. GRW9 were detected in distantly related bird species suggesting that, although each of these lineages was only detected in two bird species, they are not strict specialist parasites [32]. Additionally, *Haemoproteus* spp. PADOM05, DELURB2 lineages and *Plasmodium* spp. DELURB5, PADOM1 spp. lineages were detected in closely related bird species suggesting that these lineages may tent to be specialist parasites [32]. The prevalence displayed by these parasite lineages varied depending on the bird species infecting. Thus, in house sparrows (*Passer domesticus)* prevalence was higher for specialist parasites (*Haemoproteus* spp. PADOM05) than for generalist parasites. In contrast, Spanish house sparrows (*Passer hispaniolensis*) displayed prevalence was higher for generalist parasites (*P. relictum* SGS1). These results agree with previous studies suggesting that, in certain cases, specialist parasites might arise higher prevalence than generalist parasites [73] but, in other cases, generalist manage to reach high prevalence [70]. Our results can only confirm that depending on the parasite lineage and the host bird species prevalence might differ, suggesting that is the host-parasite interaction ultimately the factor that might affect prevalence in a wild community.

In conclusion, we have shown that, at the community level, the interaction between host and parasite identity might more strongly affect observed variation in prevalence than bird host species or parasite lineage alone. These results highlight the importance of studies focusing on the community level and analysing how different parasite lineages might interact with a variety of host species. The present study also highlights that even a parasite (such *as P. relictum* SGS1 and *H. passeris*. PADOM05) that are considered as generalist or a specialist can display both high and low prevalence per host species. Taken as a whole, these results suggest that prevalence mainly results from the interaction between host immune defences and parasite exploitation strategy where parasite lineage might play an important role following an approach depending on the bird species.

## ACKNOWLEDGMENTS

We would like to thank Sergio Magallanes for stimulating discussion and comments on this paper. We are also grateful to Vincenzo Ellis and Beatriz Willink for statistical advice and comments that greatly improved the manuscript.

